# Optimizing broadly neutralizing antibodies via all-atom interaction modeling and pre-trained language models

**DOI:** 10.64898/2026.01.20.700456

**Authors:** Yidong Song, Fandi Wu, Rubo Wang, Wei Zheng, Bing He, Qihong Yan, Xiaohan Huang, Yuquan Li, Sheng Chen, Qianmu Yuan, Jiahua Rao, Zhenchao Tang, Jiale Zhou, Haohuai He, Jincun Zhao, Yuedong Yang, Jianhua Yao

## Abstract

Antibody optimization is a fundamental challenge, and the identification of antibody-antigen interactions is crucial in the optimization process. However, current methods cannot accurately predict antibody–antigen interactions, providing limited functional guidance to improve the time-consuming and costly traditional optimization techniques. Here, we present InterAb and InterAb-Opt, a unified computational framework that integrates all-atom modeling with antibody language models to predict antibody–antigen interactions and enable antibody optimization. Leveraging the proposed all-atom modeling approach, AtomInter, and pre-trained antibody language models, InterAb outperforms existing methods in predicting antibody specificity and antibody-antigen binding affinity. InterAb successfully identified influenza A virus-binding antibodies from an antibody library and accurately detected high-affinity antibodies in the AIntibody competition. Empowered by the robust functional insights from InterAb, InterAb-Opt was developed to optimize broadly neutralizing antibodies. For R1-32 antibody, biolayer interferometry results reveal that 85%, 80%, 90%, and 67.5% of the 40 InterAb-Opt-optimized antibodies exhibit enhanced binding affinities to wild-type SARS-CoV-2, Lambda, BQ.1.1, and EG.5.1, respectively, with a maximum improvement of up to 96-fold. For the newly emerging BA.2.86 and KP.3, 55% and 52.5% of the optimized antibodies notably transition from non-binding to binding. Neutralization assays demonstrated that the optimized antibodies exhibited enhanced neutralization activity across multiple targets, highlighting the capability of InterAb-Opt in engineering broadly neutralizing antibodies. This technology enables precise analysis of antibody-antigen interactions and optimization of broadly neutralizing antibodies, holding promise for addressing challenges in immune evasion and vaccine design.

## 1 Introduction

Antibodies play an important role in the immune system by neutralizing specific pathogens [1, 2]. Optimizing the breadth of antibody neutralization against viruses is essential to address immune evasion [3, 4] and facilitate vaccine design [5, 6]. However, traditional methods for optimizing broadly neutralizing antibodies are time-consuming and costly, primarily because of the complex process of identifying antibody-antigen interactions [7–9]. Therefore, it is imperative to develop a computational method for accurately predicting antibody-antigen interactions and further promoting the optimization of broadly neutralizing antibodies.

For antibody-antigen interaction prediction, current computational methods are mainly classified into two categories according to the type of data employed, namely sequence-based methods [10, 11] and structure-based methods [12, 13]. The sequence-based methods extract features from protein sequences to determine the binding specificity and affinity. For instance, PIPR [10] utilized a deep residual recurrent convolutional neural network to extract local and contextual features from protein sequences, thereby facilitating the prediction of protein–protein interactions. Utilizing pre-trained protein-BERT [14] and AAindex [15], MVSF-AB [16] integrated semantic features and residue features to predict the binding affinity between antibodies and antigens. A2binder [17], another sequence-based method, employed sequence embeddings derived from pre-trained language models (PLMs) to identify the binding specificity and affinity of paired antibodies and antigens. Sequence-based methods exhibit low complexity and are adaptable to high-throughput screening. However, their performance is constrained by the absence of the crucial features implied by the structures, as antibody-antigen binding depends on spatial complementarity (e.g., shape matching, hydrogen bonds, and hydrophobic interactions) that cannot be fully inferred from sequence alone [18].

To incorporate structural information, structure-based methods predict the antibody-antigen interactions based on complex structures. Using graph-based signatures derived from structures, CSM-AB [12] captured close-contact information and surrounding structural features to predict the binding affinity between antibodies and antigens. AREA-AFFINITY [13] employed surface and interface areas of complex structures, including antibody surface area, antigen surface area, and interface area, to assess the antibody-antigen interactions. Despite their superior performance, the application of structure-based methods is significantly restricted by the scarcity of experimentally determined structures and their inability to predict novel proteins. In real-world applications, the interactions between antibodies and antigens (specificity and affinity) are often established prior to structural resolution, rendering methods that rely on experimentally determined complex structures less applicable in practical scenarios. Moreover, existing structure-based methods generally employ coarse-grained modeling for complex structures, resulting in an inadequate utilization of structural information, particularly the interactions between side chains.

Leveraging recent breakthroughs [19–23] in protein structure prediction, predicted structures can be used to overcome the dependence on experimentally determined structures. Addressing the limitations in structural data, all-atom modeling can further facilitate a fine-grained analysis of the interactions between antibodies and antigens. With reference to AlphaFold 3 [21], all-atom modeling was employed to accurately model the atomic-level structures of biomolecular complexes. By integrating various atomic features, an atom transformer was utilized to learn per-atom representations, which are crucial for analyzing interactions within complex structures. Beyond complex structures, the modeling of monomer structures is also important, a factor that many current methods tend to neglect. Geometric graph learning [24] holds promise for modeling monomer structures, exhibiting superior performance in enzyme function prediction [25], nucleic-acid-binding site identification [26], and protein design [27, 28]. Apart from structural information, sequence information obtained from PLMs can enhance model performance, supporting applications in drug discovery [28–30], chemical research [31], and protein intrinsic disorder prediction [32]. Considering the unique characteristics of antibody data, pre-trained models designed specifically for antibody sequences are expected to outperform other general models. By integrating the aforementioned multi-scale information, we can accurately predict antibody-antigen interactions, thereby guiding the optimization of broadly neutralizing antibodies.

In this work, we proposed InterAb and InterAb-Opt, a unified computational framework that combines all-atom modeling with antibody language models to predict antibody–antigen interactions and enable antibody optimization. In InterAb, we developed an all-atom modeling approach, AtomInter, to extract fine-grained features from complexes and innovatively apply it to analyze antibody-antigen interaction. To enhance structural representation, geometric graph learning is employed to extract geometric features from monomer structures. In addition to structural information, the antibody language models were pre-trained on over one billion antibody heavy and light chain sequences to extract comprehensive sequence embeddings. *In silico* experiments demonstrated the superior performance of InterAb in tasks related to specificity and affinity. In the antibody library we constructed, InterAb successfully identified antibodies capable of binding to influenza A virus, demonstrating its effectiveness in virtual screening. Based on InterAb, we further developed an antibody optimization framework named InterAb-Opt, aiming to address the high costs associated with traditional antibody optimization methods. In this framework, critical sites were identified using alanine scanning via InterAb, and optimized sequences were generated through our previously developed IgGM to construct a candidate antibody library. Subsequently, the library was filtered using InterAb and the sequences with higher affinity were validated by biolayer interferometry (BLI). A real-world antibody R1-32 was optimized for broad-spectrum activity against wild-type SARS-CoV-2, Lambda, BQ.1.1, EG.5.1, BA.2.86, and KP.3. Among the 40 optimized antibodies, 85%, 80%, 90%, and 67.5% exhibited enhanced binding affinities to wild-type SARS-CoV-2, Lambda, BQ.1.1, and EG.5.1, respectively, with the highest improvement exceeding 96-fold. A total of 22 and 21 optimized antibodies achieved a remarkable transition from non-binding to binding for BA.2.86 and KP.3. Neutralization assays revealed that 85% of the optimized antibodies showed improved neutralizing activity against at least one antigen, demonstrating the outstanding capability of InterAb-Opt in the optimization of broadly neutralizing antibodies.

## 2 Results

### 2.1 Interaction-guided antibody optimization

The specific binding (Fig. 1a) between antibodies and antigens represents a critical interaction within the immune system, and elucidating this interaction is essential for antibody optimization. Here, we propose a computational method, termed InterAb-Opt, to optimize the breadth of antibodies (Fig. 1b) through all-atom interaction modeling. InterAb-Opt generates candidate sequences via a generative model, and employs our proposed InterAb framework to identify critical sites and screen for optimized antibodies(Fig. 4a). InterAb is an all-atom-based method designed to predict antibody specificity and antibody-antigen binding affinity, and has been further applied to enable the optimization of broadly neutralizing antibodies. The model integrates sequence information, monomer structures, and complex structures (Fig. 1c), while incorporating both inner-chain and inter-chain information. Monomer and complex structures were predicted using ESMFold and Chai-1, respectively. InterAb primarily consists of three modules: the Inner-chain sequence module, the Inner-chain structure module, and the Inter-chain module. The Inner-chain sequence module utilizes pre-trained antibody language models [17] and ESM2 [23] to extract sequence embeddings from antibody and antigen sequences (Fig. 1d). The Inner-chain structure module acquires monomer embeddings via geometric graph learning, relying on the monomer structures of antibodies and antigens (Fig. 1e). The Inter-chain module introduces AtomInter, an all-atom modeling approach that extracts complex embeddings from antibody–antigen complexes (Fig. 1f). The AtomInter first extracts the antibody-antigen interface from complexes based on interatomic distances and generates atom-level features, then employs an atom transformer to learn the interactions between the antibody and antigen. The embeddings of the Inner-chain sequence module, Inner-chain structure module, and Inter-chain module are integrated as multi-scale information to identify the binding specificity and affinity between antibodies and antigens.

**Fig. 1.**
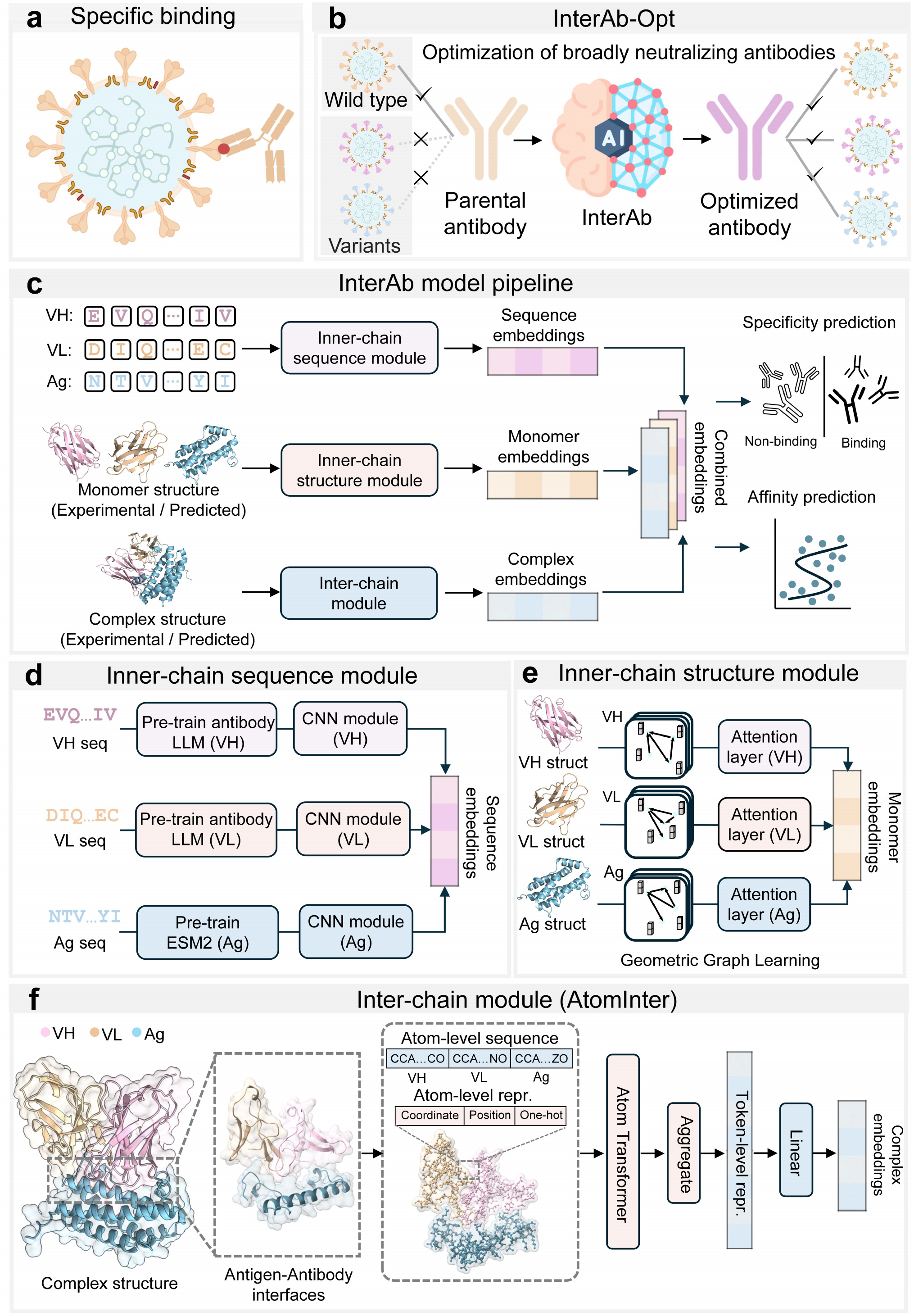
The architecture of InterAb. (**a**) Schematic illustration of the specific binding between an antibody and an antigen. (**b**) Broadly neutralizing antibody optimization using InterAb-Opt. The parental antibody was unable to bind antigen variants, but following optimization, it gained the capacity to recognize and bind multiple antigen variants, thereby achieving broad-spectrum reactivity. (**c**) The binding specificity and affinity between antibodies and antigens were identified by InterAb. Given the input sequences, the Inner-chain sequence module utilizes pre-trained antibody language models and ESM2 to generate sequence embeddings. To extract features of monomer structures, the monomer embeddings were derived by Inner-chain structure module using ESMFold-predicted structures of antibodies and antigens. For the enhanced capture of inter-chain interactions, the Inter-chain module is employed to extract complex embeddings from the antibody-antigen complex structures. The sequence embeddings, monomer embeddings, and complex embeddings were integrated to form combined embeddings, which were subsequently utilized for the prediction of specificity and affinity. (**d**) Inner-chain sequence module. Pre-trained antibody language models were utilized to extract sequence features from the heavy and light chains of antibodies, while the ESM2 model was employed to derive evolutionary information from antigen sequences, collectively forming the sequence embeddings. (**e**) Inner-chain structure module. Based on the monomer structures of antibodies and antigens, geometric graph learning was employed to derive monomer embeddings. (**f**) Inter-chain module. AtomInter was proposed for all-atom modeling of antibody-antigen complexes. From antibody-antigen complex structures, the interfaces between antibodies and antigens were extracted, followed by the derivation of the atom-level sequences. The atom-level sequence is encoded via one-hot encoding and integrated with coordinate and position information to form the atom-level representations. These representations are then processed through an atom transformer to generate complex embeddings.

### 2.2 InterAb accurately predicts antibody specificity

InterAb was evaluated for antibody specificity prediction utilizing the specificity dataset SPE7620 (details shown in Section 4.1). Antibody specificity prediction refers to the task of predicting whether a given antibody binds to a target antigen. Fig. 2a presents a comparative analysis of the receiver operating characteristic curves for InterAb and five other methods (A2binder [17], AbAgIntPre [33], AntiBinder [34], S3AI [11], and PIPR [10]) on SPE-Test. InterAb achieved the highest AUC (area under the receiver operating characteristic curve) value of 0.9295, surpassing the second-best method A2binder (AUC=0.8878) by 4.7%, thus demonstrating the efficacy of InterAb. A2binder derived comprehensive sequence features from pre-trained language models, yet it failed to extract structural information from the protein structures. AbAgIntPre adopts a siamese-like convolutional neural network architecture, achieving the third-ranked performance with an AUC value of 0.8812. Supplementary Fig. S1 displays the precision-recall curves of these methods, with InterAb achieving an AUPR (area under the precision-recall curve) of 0.8845, surpassing A2binder (AUPR=0.8210) and AbAgIntPre(AUPR=0.7393) by 7.7% and 19.6%, respectively.

**Fig. 2.**
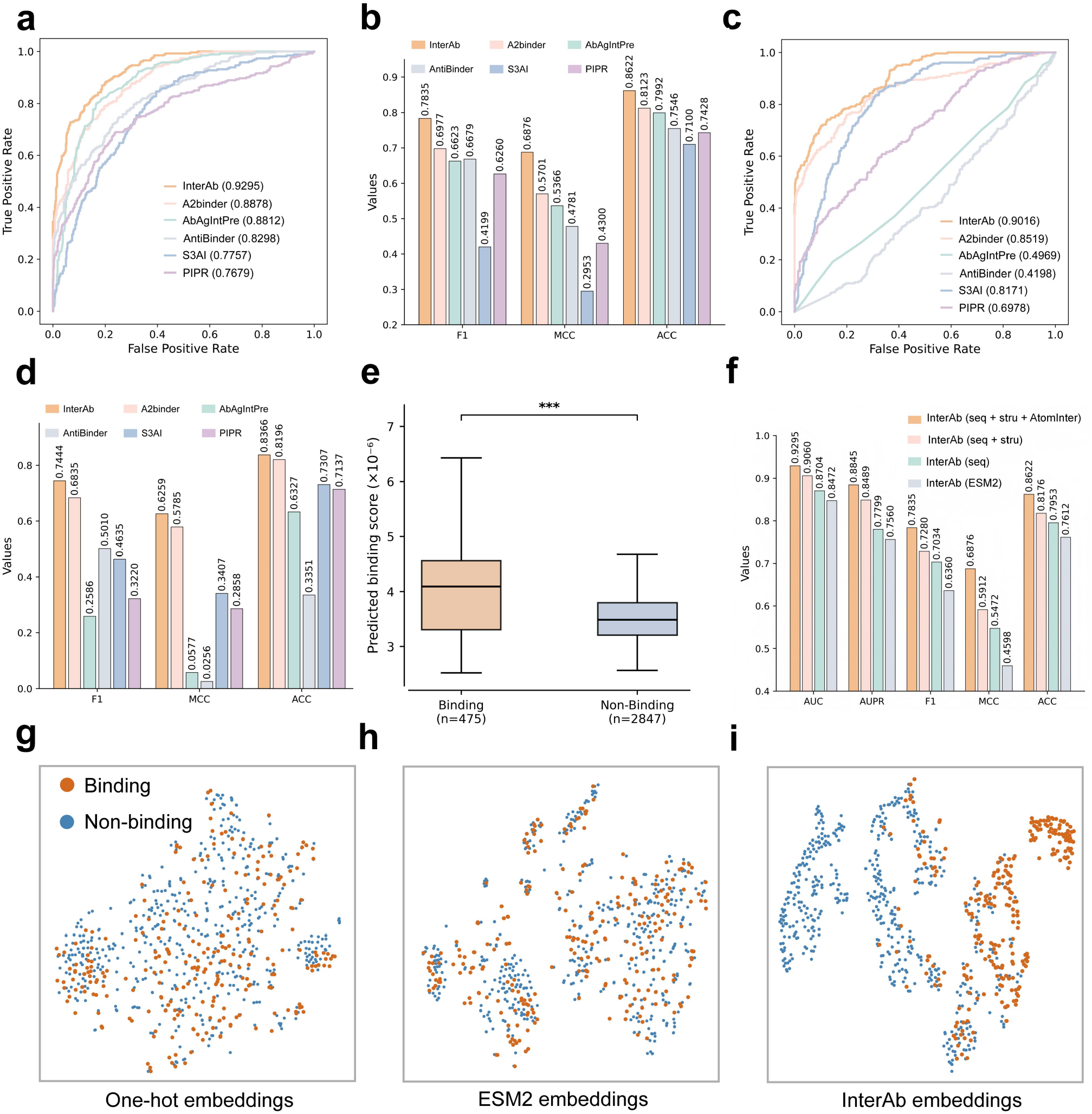
Antibody specificity prediction. (**a**) The receiver operating characteristic curves of InterAb and five comparative methods on SPE-Test. (**b**) InterAb was evaluated with A2binder, AbAgIntPre, AntiBinder, S3AI, and PIPR using F1, MCC, and ACC on the SPE-Test dataset. (**c**) The receiver operating characteristic curves of InterAb on the unseen test set, in comparison with five previous methods. (**d**) A comparative analysis of InterAb and five other methods was conducted on unseen data, with evaluation metrics including F1, MCC, and ACC. (**e**) Comparison of predicted scores of InterAb between binding and non-binding antibodies in the phage display library targeting H3N2. (**f**) An ablation study was conducted to evaluate the contributions of the Inner-chain sequence module (seq), Inner-chain structure module (stru), and Inter-chain module (AtomInter) to the performance of the model. The InterAb (ESM2) denotes the replacement of the antibody language models in the Inner-chain sequence module with ESM2. InterAb (seq) refers to the use of only the Inner-chain sequence module. InterAb (seq+stru) indicates the utilization of both Inner-chain sequence module and Inner-chain structure module. InterAb (seq+stru+AtomInter) represents the use of Inner-chain sequence module, Inner-chain structure module, and AtomInter in Inter-chain module. (**g, h, i**) The discriminative capabilities of one-hot embeddings, ESM2 embeddings, and InterAb embeddings were evaluated for distinguishing between binding and non-binding antibody-antigen pairs.

To further evaluate the model performance, additional metrics such as F1, Matthews correlation coefficient (MCC), and Accuracy (ACC) were employed (Fig. 2b). InterAb demonstrated superior results across all three metrics, achieving an F1 of 0.7835, an MCC of 0.6876, and an ACC of 0.8622. These values exceed those of the second-best method (A2binder) by 12.3%, 20.6%, and 6.1%. We then evaluated the performance of the model on unseen antigens. Similarly, we randomly divided SPE7620 into training (80%), validation (10%), and test sets (10%), while ensuring that antigens in the test set were excluded from the training and validation sets. The performance of InterAb, along with five competing methods, was rigorously assessed under the more challenging setting of unseen data.

Fig. 2c presents the receiver operating characteristic curves of these methods. InterAb achieved an AUC of 0.9016, outperforming A2binder (0.8519) and demonstrating its advantage on the unseen test set. As shown in Fig. 2d, InterAb achieved the highest performance with an F1 of 0.7444, an MCC of 0.6259, and an ACC of 0.8366. The performance of A2binder was significantly constrained by its dependence on homologous sequence information, resulting in F1, MCC, and ACC values lower by 8.2%, 7.6%, and 2.0% than those of InterAb. For unseen antigen data, AbAgIntPre and AntiBinder showed a substantial performance drop, with F1 scores of 0.2586 and 0.5010, respectively, highlighting the inherent challenge of this setting. Comparisons of precision-recall curves across all methods are provided in Supplementary Fig. S2. Evaluation on unseen data confirms that InterAb consistently exhibits high predictive accuracy for novel antigens, highlighting its substantial potential in the field of drug discovery.

### 2.3 InterAb can identify influenza A-specific antibodies

In our previous work on the tFold System [35], we constructed a phage-displayed antibody library specifically targeting influenza A (H3N2). To identify antibodies within the library capable of binding to H3N2, individual phage clones were screened via ELISA. His-tagged protein served as the negative control, and clones with an OD450 value greater than three times that of the negative control were defined as positive binders. The positive clones were subsequently subjected to Sanger sequencing to confirm their sequences, yielding a total of 475 unique sequences. Given that ELISA-based validation is both time-consuming and costly, we herein explore the use of InterAb to accelerate this screening process.

InterAb was employed to identify antibodies from the library that demonstrated specific binding to H3N2 via ELISA validation. A dataset was constructed by designating ELISA-confirmed binders as positive samples (475) and selecting antibodies from the library with CDR-H3 and CDR-L3 similarity below 0.7 to the positive samples as negative controls (2847). InterAb was fine-tuned on FluA-Data and subsequently evaluated on this custom dataset. Fig. 2e presents the scoring results of the dataset using InterAb. ELISA-confirmed binders received significantly higher prediction scores from the model compared to non-binders (*p* < 0.001), with an average score exceeding that of non-binders by 13.2%. The enrichment factor at the 1% screening rate (EF1%) reached 6.57, demonstrating the capacity of InterAb to effectively enrich influenza A-specific binders from antibody library.

### 2.4 Ablation experiments on InterAb

Ablation studies were performed to analyze the contributions of different modules. Fig. 2f illustrates the impact of the Inner-chain sequence module, Inner-chain structure module, and Inter-chain module on the model performance. Using only the Inner-chain sequence module, the model achieved an AUC of 0.8704 and an AUPR of 0.7799 on SPE-Test. By leveraging the residue-level information derived from pre-trained antibody language models, the Inner-chain sequence module achieved acceptable results. When the antibody language models in the Inner-chain sequence module were replaced with ESM2, the model’s AUC and AUPR decreased to 0.8472 and 0.7560, respectively, demonstrating the effectiveness of antibody language models. The inclusion of the Inner-chain structure module led to enhancements of 4.1% in AUC and 8.8% in AUPR, demonstrating that geometric graph learning can learn effective features from monomer structures. When adding the AtomInter of Inter-chain module, the model performance was further improved, with AUC and AUPR values increasing by 2.6% and 4.2%, respectively, validating the effectiveness of all-atom modeling for complex structures. For other metrics, the addition of the AtomInter also has the best results, with values of 0.7835, 0.6876, and 0.8622 for F1, MCC, and ACC, respectively. These results demonstrate that all-atom modeling enables the extraction of critical interaction information from complex structures.

To elucidate the exceptional performance, we analyzed the t-SNE projection of the embeddings learned by the model. As illustrated in Fig. 2g, the one-hot embeddings obtained by encoding the sequence in a one-hot manner failed to differentiate between binding and non-binding data. Despite the improved results generated by ESM2 embeddings (Fig. 2h), the points representing both binding and non-binding instances persisted in a mixed distribution. The embeddings learned through InterAb effectively distinguished between these two distinct types of data (Fig. 2i), indicating that InterAb can accurately identify antibody specificity.

### 2.5 InterAb outperforms existing methods in affinity prediction

Antibody-antigen binding affinity prediction presents greater challenges than specificity prediction, yet InterAb achieved robust performance in this task. Affinity prediction refers to the task of predicting the binding affinity of a given antibody-antigen pair. We evaluated the performance of InterAb and four other methods (A2binder, PIPR, ESM2, and CSM-AB) in the antibody–antigen binding affinity task. ESM2 denotes the use of the ESM2 model to encode antibody and antigen sequences, with the resulting feature representations employed for affinity prediction. As shown in Fig. 3a, InterAb achieved a Pearson’s correlation of 0.6580 on AFF-Test, surpassing A2binder (Pearson’s r = 0.6190, Fig. 3b) and PIPR (Pearson’s r = 0.5914, Fig. 3c) by 6.3% and 11.3%, respectively. This indicates the effectiveness of InterAb in predicting antibody–antigen binding affinity. For Spearman’s correlation, RMSE (Root mean square error), and MAE (Mean absolute error), InterAb also attained the best results of 0.6520, 1.9151, and 1.4202 (Fig. 3, d, e, f), respectively. The sequence-based method A2binder achieved the second-best performance by learning internal sequence features, with Spearman’s correlation, RMSE, and MAE values of 0.5976, 2.3032, and 1.6688, respectively. CSM-AB, a graph-based method, achieved acceptable results by modeling interaction interfaces as graph-based signatures, yielding Spearman’s correlation, RMSE, and MAE values of 0.1294, 2.6400, and 1.9726, respectively. Unlike CSM-AB, InterAb employed all-atom geometric representations instead of coarse-grained graph features, allowing it to capture fine-grained interactions between antibodies and antigens. We further compared the results obtained using experimentally determined structures in AtomInter. Under this condition, the model achieved superior performance, with Pearson’s and Spearman’s correlation coefficients of 0.6877 and 0.6751, respectively (Supplementary Fig. S4 and Table S7). These results demonstrate that the model attains better performance when experimental structures are available.

**Fig. 3.**
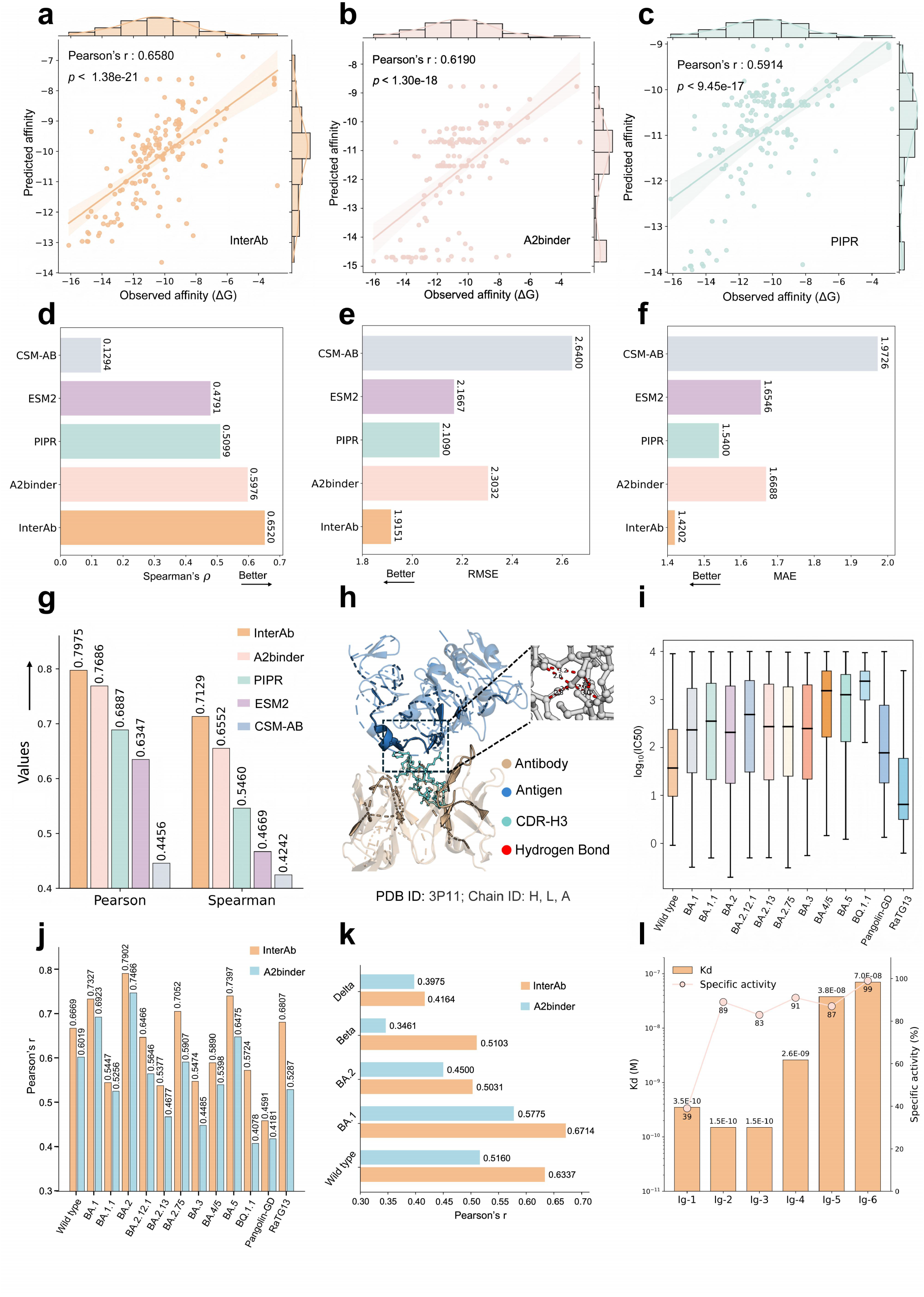
Antibody–antigen binding affinity prediction. (**a**) The scatter plot illustrates the correlation between experimentally validated and InterAb-predicted binding affinity values on AFF-Test, with similar comparisons made for A2binder (**b**) and PIPR (**c**). (**d**) The Spearman’s correlation of InterAb was compared with those from four other methods (A2binder, PIPR, ESM2, and CSM-AB) on AFF-Test. (**e**) The Root mean square error (RMSE) and Mean absolute error (MAE) (**f**) were analyzed between InterAb and other methods. (**g**) The performance of InterAb and comparative methods was evaluated on SKE-Test using Pearson’s and Spearman’s correlation coefficients. (**h**) The attention scores of the model were visualized for one example (3P11), with highlighted portion representing higher attention scores. The model exhibited higher attention scores in the CDR-H3 regions of antibodies, while regions on the antigen with high attention scores corresponded to locations of hydrogen bonds. This demonstrates the model’s ability to focus on critical regions of both antibodies and antigens. (**i**) The distribution of the log-transformed half maximal inhibitory concentration (*log*_10_(*IC*_50_)) values for the datasets corresponding to the wild-type SARS-CoV-2, its ten variants, and two related coronaviruses (Pangolin-GD and RaTG13) was visualized. (**j**) The Pearson’s correlation between the predicted and observed *log*_10_(*IC*_50_) values were analyzed for both InterAb and A2binder across multiple datasets, including the wild-type SARS-CoV-2, its ten variants, as well as the related coronaviruses Pangolin-GD and RaTG13. (**k**) The Pearson’s correlation between the predicted and observed *pK*_*d*_ values for InterAb and A2binder was analyzed on the wild-type SARS-CoV-2 dataset and four additional variant datasets. (**l**) The *K*_*d*_ values and specific activities of antibodies optimized in the AIntibody competition.

To conduct a comprehensive performance evaluation, we established a new dataset SKE426 from the SKEMPI 2.0 [36] database and applied it to test different methods. The dataset was labeled with the log-transformed dissociation constant (*pK*_*d*_), with 80% of the data randomly allocated to the training set, 10% to the validation set, and the remaining 10% to the test set (SKE-Test). Fig. 3g illustrates the correlation between experimentally validated *pK*_*d*_ values and those predicted by InterAb and other competing methods on SKE-Test. InterAb achieves Pearson’s and Spearman’s correlation coefficients of 0.7975 and 0.7129, outperforming A2binder by 3.8% and 8.8%, respectively. In comparison with four other methods, InterAb exhibited superior performance, attaining an RMSE of 0.8009 and an MAE of 0.6106 (Supplementary Fig. S8). The RMSE values for A2binder, PIPR, and ESM2 were 0.8610, 1.1283, and 1.0252, respectively, which are higher than that of InterAb by 7.5%, 40.9%, and 28.0%. CSM-AB exhibited high error on the SKE-Test (RMSE = 1.3942, MAE = 1.1114), likely due to its reliance on coarse-grained graph topology, which cannot capture atomic-level spatial changes induced by mutations. These results further substantiate the efficacy of InterAb in predicting antibody-antigen binding affinity.

### 2.6 All-atom modeling enables accurate affinity prediction

We investigated the effect of all-atom modeling implemented through AtomInter in the Inter-chain module. When AtomInter was not utilized, the Pearson’s correlation of the model using only sequence information was 0.5904 (Supplementary Fig. S9), and it reached 0.6311 after incorporating monomer structure information. When integrating AtomInter for all-atom modeling, the performance of the model significantly improved, with the Pearson’s correlation reaching 0.6580. This finding demonstrates that the application of AtomInter can accurately predict the interactions between antibodies and antigens. We further ran t-SNE to analyze the embeddings learned by InterAb. As illustrated in Supplementary Fig. S11, the ESM2 embeddings failed to distinguish between points with binding free energy (Delta G) < -10 kcal/mol and those with Delta G ≥ -10 kcal/mol. Delta G ≥ -10 kcal/mol indicates a weaker binding affinity, whereas Delta G < -10 kcal/mol suggests a stronger binding affinity [37, 38]. In contrast, InterAb demonstrated superior capability in distinguishing between data points with these two distinct levels of binding affinity. These results provide additional insights into the exceptional performance of InterAb and further demonstrate the effectiveness of AtomInter.

We further analyzed the attention scores extracted from the model’s attention layer and found that InterAb can effectively focus on critical regions of antibody-antigen interactions. Fig. 3h illustrates a case (3P11 [39]), where darker colors indicate higher attention scores, reflecting enhanced focus on this region. In the antibody portion, the model exhibited higher attention scores in the CDR-H3 regions, consistent with the essential role these regions play in mediating antibody-antigen interactions. In the antigen portion, the model demonstrated higher attention scores at the interface with the antibody. Further analysis revealed that regions on the antigen with high attention scores contain hydrogen bonds that are important for the interaction. These results indicate that InterAb is capable of capturing critical regions involved in the antibody-antigen interaction. For additional examples, please refer to Supplementary Figs. S12 and S13.

### 2.7 InterAb demonstrates adaptability to SARS-CoV-2

By integrating our experimental data with publicly available datasets, we constructed a comprehensive and up-to-date dataset for SARS-CoV-2. Fig. 3i displays the distribution of data annotated with log-transformed half maximal inhibitory concentration (*log*_10_*IC*_50_), including the wild type and 10 variants of SARS-CoV-2, as well as two closely related coronaviruses (Pangolin-GD and RaTG13). Compared to the wild type, the binding affinity between other variants and antibodies gradually decreases, reflecting the immune escape effect of mutant strains. For each antigen, we randomly allocated 80% of the data for training, 10% for validation, and utilized the remaining 10% as the test set. In these data, InterAb exhibited robust Pearson’s correlation (Fig. 3j), with Pearson’s correlation exceeding 0.6 for over 53.8% of the antigens, thereby demonstrating its superior predictive capability across different variant strains. Supplementary Figs. S14 to S16 shows the correlation between the observed *log*_10_*IC*_50_ values and the *log*_10_*IC*_50_ values predicted by InterAb across the datasets for wild-type SARS-CoV-2 (Spearman’s *ρ* = 0.6764), BA.1 (Spearman’s *ρ* = 0.7310), and BA.2 (Spearman’s *ρ* = 0.7830). Results for other antigens are presented in Supplementary Fig. S17. Additional comparisons of RMSE and MAE are available in Supplementary Table S8. The aforementioned results reveal that InterAb is capable of accurately predicting the binding affinity between different mutant viruses and antibodies.

In addition to the data annotated with *IC*_50_, we have also curated data annotated with dissociation constant (*K*_*d*_). The *K*_*d*_ values were further transformed into the logarithmic space as *pK*_*d*_, and Supplementary Fig. S19 illustrates the distribution of the *pK*_*d*_ values for the wild type and four variants (BA.1, BA.2, Beta, Delta). Through logarithmic transformation, the *pK*_*d*_ values are primarily concentrated between 8 and 10. Fig. 3k demonstrates that InterAb achieved a Pearson’s correlation of 0.6337 on the wild-type dataset, surpassing A2binder by 22.8%. InterAb also demonstrated strong performance across other variant strains, with Pearson’s correlation of 0.6714, 0.5031, 0.5103, and 0.4164 for BA.1, BA.2, Beta, and Delta, respectively. Supplementary Fig. S20 presents scatter plots comparing the performance of InterAb and the second-best method, A2binder, on the wild-type dataset. Additional results regarding RMSE and MAE are provided in Supplementary Table S9. These consistent results provide robust evidence for InterAb’s capability to predict interactions between various SARS-CoV-2 variants and antibodies.

In the AIntibody competition [40] reported in Nature Biotechnology, SARS-CoV-2-targeting antibodies predicted by InterAb exhibited outstanding binding capacity and functional activity. Fig. 3l presents the *K*_*d*_ values of six submitted antibodies measured by surface plasmon resonance (SPR), among which 67% exhibited affinity at the nanomolar level, with the highest reaching 150 pM. Of the submitted antibodies, over 83% demonstrated specific activity exceeding 80% (Fig. 3l), with the maximum reaching 99%, indicating strong activity. Supplementary Fig. S21 illustrates the average association rate constants of the antibodies with their respective antigens, revealing that Ig-1, Ig-2, and Ig-3 achieved high affinity through rapid binding kinetics, underscoring InterAb’s capacity to capture mechanistic insights at the kinetic level. The core functionality underlying the practical utility of the designed antibodies was further validated via baculovirus particle (BVP) enzyme-linked immunosorbent assay (ELISA) (Supplementary Fig. S21). Collectively, these results substantiate InterAb’s robust capability in guiding wet-lab experiments and underscore its substantial value in antibody optimization.

### 2.8 Applying InterAb-Opt for the optimization of broadly neutralizing antibodies

Building on the functional insights derived from InterAb, we developed InterAb-Opt (Fig. 4a), a framework for broadly neutralizing antibody optimization. InterAb-Opt initially identified critical sites on the original antibody based on distance and alanine scanning using InterAb. Subsequently, a candidate antibody library was constructed using our previously developed IgGM [30]. The candidate antibody library was then screened with InterAb to identify optimized antibodies with broad-spectrum activity, which were further validated through wet-lab experiments using BLI. A real-world antibody, R1-32 [41], was optimized using InterAb-Opt to validate its effectiveness. The R1-32 antibody targets the SARS-CoV-2 receptor-binding domain (RBD) with nanomolar affinity *K*_*d*_ =1.35 nM, potently neutralizing wild-type SARS-CoV-2 by inducing conformational changes in the RBD and disrupting spike protein stability. However, significant immune escape by SARS-CoV-2 variants has been observed. For Lambda, BQ.1.1, and EG.5.1, the affinity (*K*_*d*_) between R1-32 and these variants decreases to 1090 nM, 443 nM, and 60.5 nM, respectively, leading to a reduction or even loss of the original efficacy of the antibody. More concerningly, the R1-32 antibody fails to bind to the newly emerging variants BA.2.86, and KP.3. To overcome the immune evasion, InterAb-Opt was employed to optimize R1-32 with broad-spectrum activity, addressing the challenge of traditional wet-lab methods, which are often characterized by complex procedures, extended timelines, and high costs.

**Fig. 4.**
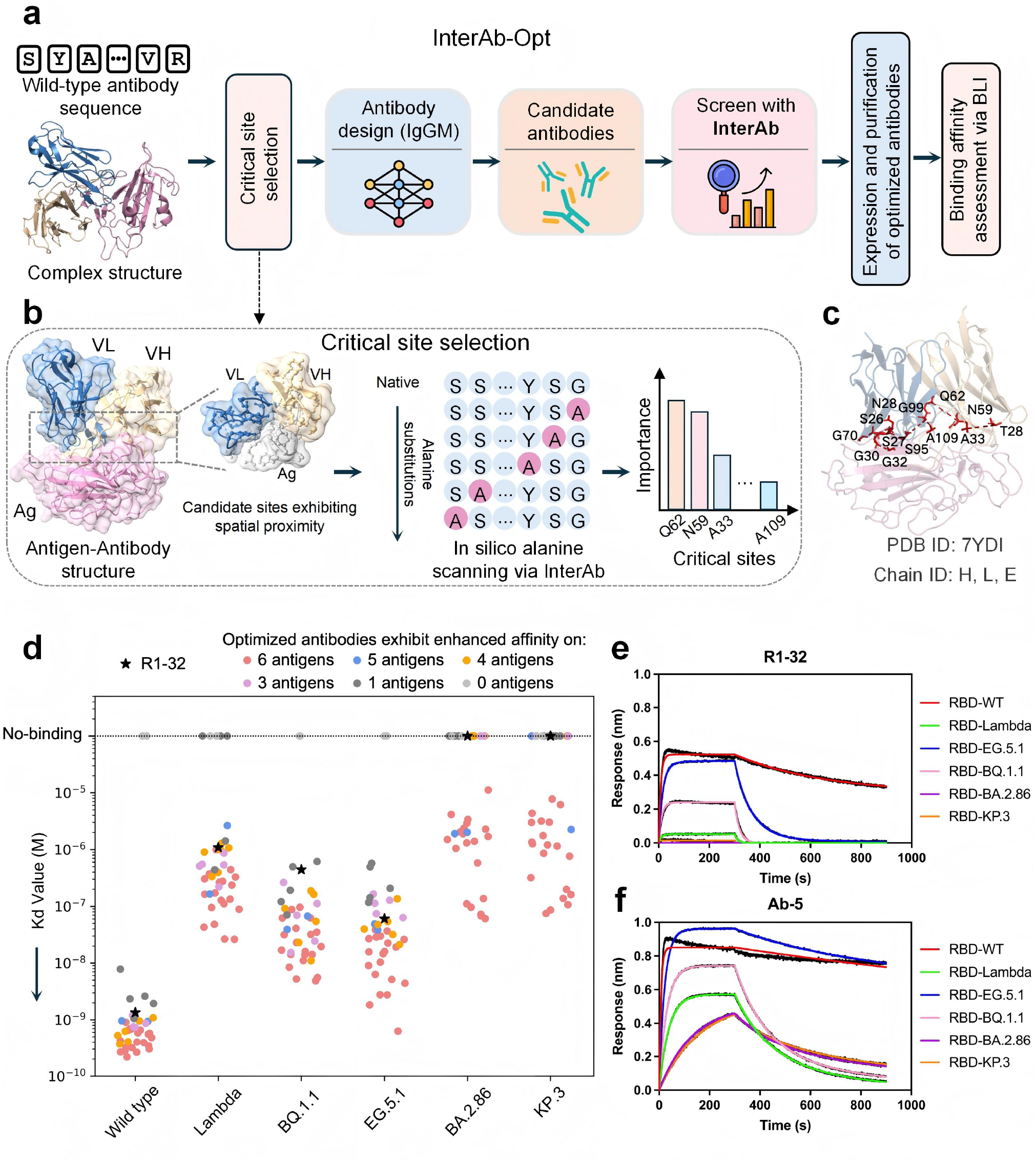
Broadly neutralizing antibodies were optimized using InterAb-Opt. (**a**) The workflow for antibody optimization using InterAb-Opt encompasses the selection of critical sites, the generation of candidate antibodies, and the screening of broadly neutralizing antibodies through InterAb. (**b**) The selection process of critical sites. Based on the antibody-antigen complex, the distances between the *C*_*α*_ atoms of residues on the antibody and those on the antigen are calculated, with residues exhibiting distances less than 8 Å being selected as candidate sites. Subsequently, *in silico* alanine scanning via InterAb is employed to determine the critical sites based on these candidate sites. (**c**) Visualization of the 13 selected critical sites was performed. (**d**) The binding affinities of the 40 optimized antibodies against wild-type SARS-CoV-2, Lambda, BQ.1.1, EG.5.1, BA.2.86, and KP.3 were evaluated and compared with those of the R1-32 antibody. (**e**) The binding curves of R1-32 and Ab-5 (**f**) were compared.

We optimized 40 novel antibodies for the wild-type SARS-CoV-2, as well as the variants Lambda, BQ.1.1, EG.5.1, BA.2.86, and KP.3 (Supplementary Table S11). For each of the six antigens, we generated 1,000 antibody sequences using IgGM and compiled them into a candidate antibody library. The affinities of candidate antibodies against all antigens were evaluated using InterAb, and the top 40 antibodies with the highest average affinities across these antigens were selected for experimental validation via biolayer interferometry (BLI) (Supplementary Table S11). Fig. 4d shows the binding affinities of the optimized antibodies to each antigen, in comparison with those of the R1-32 antibody. Among the optimized antibodies, 34 out of 40 (85%) exhibited higher binding affinities to the wild-type SARS-CoV-2 compared to R1-32, while 32 (80%), 36 (90%), and 27(67.5%) demonstrated superior affinity to the Lambda, BQ.1.1, and EG.5.1 variants, respectively (Supplementary Table S11). 22 (55%) and 21 (52.5%) optimized antibodies transitioned from non-binding to binding on the BA.2.86 and KP.3 variants. Among the antibodies, 50% (20) exhibited increased affinity across all six antigens, 95% (38) showed enhanced affinity on at least one antigen. The enhanced binding affinities across these antigens underscore the broad-spectrum activity of the optimized antibodies.

As shown in Fig. 4e,f, the binding curves of R1-32 and the optimized antibody Ab-5 against six antigens were compared. Ab-5 exhibited improved binding affinities across all six antigens compared to R1-32, with the highest affinity reaching 0.35 nM, demonstrating the effectiveness of InterAb-Opt in optimizing broadly neutralizing antibodies. In addition to Ab-5, Supplementary Fig. S22 also presents the other nine antibodies from the top ten optimized antibodies. These antibodies exhibited superior binding affinities to R1-32 across all the six antigens, thereby demonstrating their broad-spectrum activity. More results can be seen in Supplementary Figs. S23 to S24. We then analyzed the fold increase in binding affinity of the optimized antibodies relative to R1-32. Fig. 5a illustrates the fold enhancement in binding affinity of each optimized antibody against wild-type SARS-CoV-2, Lambda, BQ.1.1, and EG.5.1 compared to R1-32. Among them, 5 antibodies exhibited a more than 50-fold increase in affinity (with the highest at 96-fold), and 21 antibodies showed a more than 10-fold increase, indicating a significant improvement over R1-32. These results reveal the broad-spectrum activity of optimized antibodies across all six antigens, highlighting the superior capabilities of InterAb-Opt in broadly neutralizing antibody optimization.

**Fig. 5.**
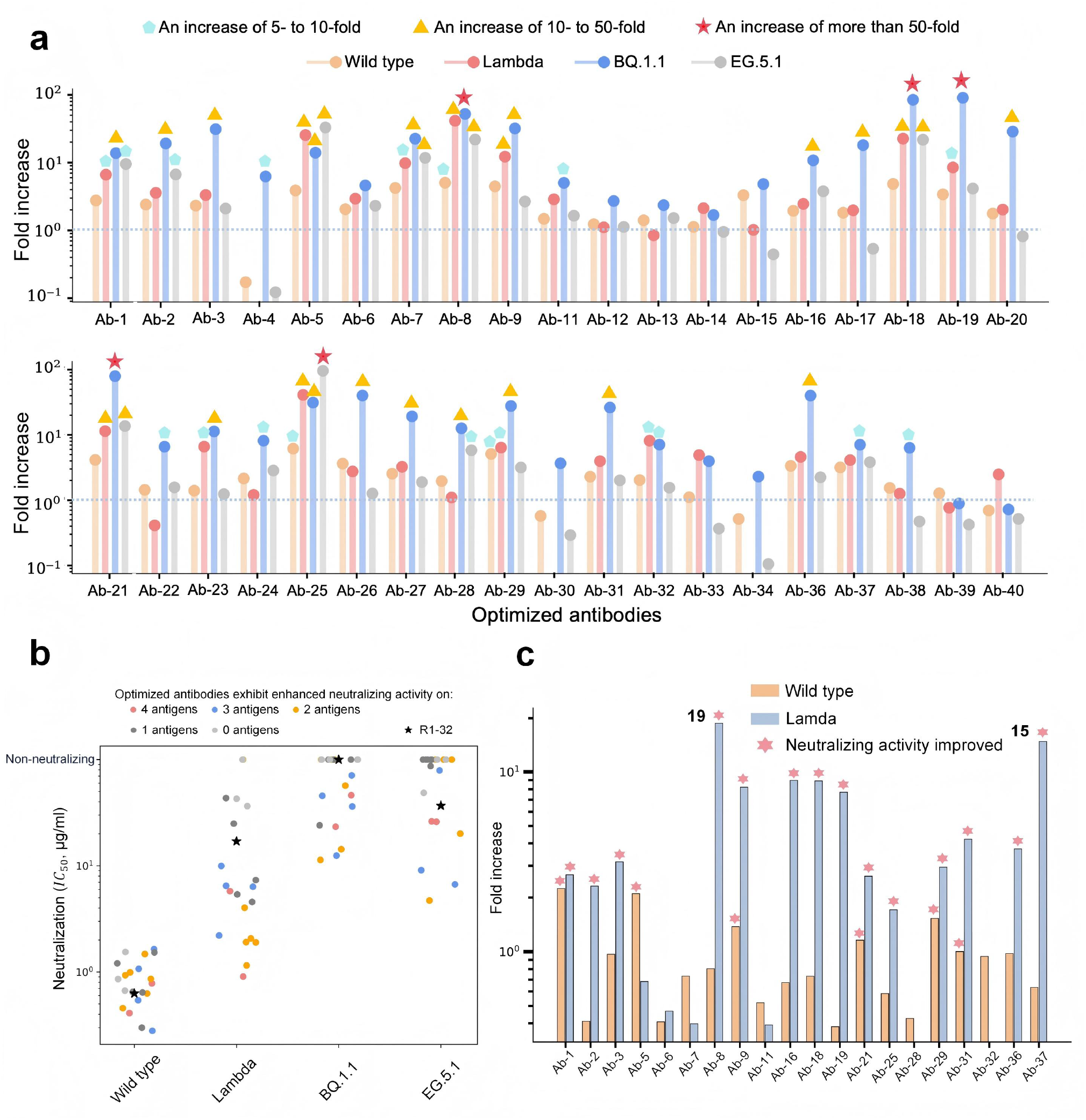
Optimized antibodies exhibit enhanced affinity and neutralizing activity. (**a**) The fold increase in binding affinity of the optimized antibodies against wild-type SARS-CoV-2, Lambda, BQ.1.1, and EG.5.1 was analyzed in comparison to the R1-32 antibody. (**b**) Neutralizing activity of the optimized antibodies against wild-type SARS-CoV-2, Lambda, BQ.1.1, and EG.5.1, compared with R1-32. (**c**) Fold increase in *IC*_50_ of optimized antibodies relative to R1-32 against wild-type SARS-CoV-2 and Lambda.

Furthermore, we assessed the neutralizing activity of the optimized antibodies by measuring the half-maximal inhibitory concentration (*IC*_50_). The top 20 antibodies with the highest affinity were selected for neutralization assays against wild-type SARS-CoV-2, Lambda, BQ.1.1, and EG.5.1. As shown in Fig. 5b, 85% of the optimized antibodies exhibited enhanced neutralization against at least one antigen, and 50% of them transitioned from non-neutralizing to neutralizing against the BQ.1.1. Fig. 5c displays the fold increase in *IC*_50_ of the optimized antibodies relative to R1-32. Due to the weak or undetectable neutralization by R1-32 against BQ.1.1 and EG.5.1, fold increase was calculated only for wild-type SARS-CoV-2 and Lambda. Among these, 75% of the optimized antibodies showed improved *IC*_50_ against at least one antigen, with a maximum improvement of 19-fold, demonstrating that the optimized antibodies exhibit substantially enhanced neutralization activity. Supplementary Fig. S25 presents the neutralization curves of the optimized antibodies compared to R1-32.

## 3 Discussion

This study presents InterAb and InterAb-Opt, a unified computational framework that combines all-atom modeling with antibody language models to characterize antibody–antigen interactions and enable antibody optimization. Leveraging the developed AtomInter, InterAb innovatively employs all-atom modeling to accurately predict antibody-antigen interactions. To improve model performance, the pre-trained antibody language models are employed to extract comprehensive antibody sequence information. Building on InterAb, InterAb-Opt is further constructed for the computational optimization of broadly neutralizing antibodies. InterAb-Opt integrates functional insights from InterAb to guide antibody optimization, overcoming the time and cost limitations of traditional techniques. Experimental validation confirmed that the model accurately identified influenza A virus-binding antibodies from an antibody library and successfully optimized the R1-32 antibody for broad neutralizing activity against six divergent antigens.

Antibody optimization [42] is essential for therapeutic antibody development, yet accurate characterization of antibody–antigen interactions represents a major bottleneck for traditional optimization approaches. Existing methods cannot accurately predict antibody–antigen interactions due to the absence of all-atom modeling, thereby failing to improve the traditional optimization techniques. In addition, these interaction prediction methods overly rely on experimentally determined structures and mostly neglect the important role of monomer structures. To address these challenges, we introduce InterAb, a novel interaction predictor that extracts fine-grained structural features via an all-atom modeling approach. To enhance structural information, the antibody language models pre-trained on over one billion antibody heavy and light chain sequences are employed to extract comprehensive sequence information. Furthermore, geometric graph learning is employed to extract geometric features of monomer structures, addressing the neglect in existing methods. Leveraging breakthroughs in structural prediction, our method eliminates reliance on experimentally determined structures, allowing for effective predictions of newly discovered antibodies. Experiments demonstrate that InterAb surpasses existing methods in predicting antibody specificity and antibody-antigen binding affinity. Furthermore, InterAb successfully discovered influenza A virus-binding antibodies from an antibody library and accurately identified high-affinity antibodies in the AIntibody competition.

Based on the functional insights from InterAb, we develope InterAb-Opt, an antibody optimization framework that successfully improves the broad-spectrum activity of R1-32 antibody [41]. InterAb-Opt is employed to optimize broad-spectrum activity for R1-32 against wild-type SARS-CoV-2, Lambda, BQ.1.1, EG.5.1, BA.2.86, and KP.3. Biolayer interferometry results reveal that 85%, 80%, 90%, and 67.5% of the 40 optimized antibodies exhibit enhanced binding affinities to wild-type SARS-CoV-2, Lambda, BQ.1.1, and EG.5.1, respectively, with the maximal improvements exceeding 96-fold. For BA.2.86 and KP.3 that do not bind to R1-32, 22 (55%) and 21 (52.5%) of the optimized antibodies achieve a remarkable transition from non-binding to binding, effectively mitigating the issue of immune escape. Neutralization assays demonstrated that 85% of the optimized antibodies exhibited enhanced neutralization against at least one antigen, with a maximum improvement of 19-fold. These results underscore the robust capability of InterAb-Opt in the optimization of broadly neutralizing antibodies.

While the model has demonstrated excellent performance, there are several areas for potential enhancement. Due to substantial storage requirements, we currently limit our all-atom modeling to the interface area of antibodies and antigens. If future storage constraints are alleviated, utilizing the entire structure for all-atom modeling could achieve superior results. Besides, because of the scarcity of antibody-antigen specific and affinity data, the model is trained on a limited dataset. Accumulating more data for training in the future could further improve the model performance.

In summary, we propose an antibody-antigen interaction predictor InterAb, and an antibody optimization framework InterAb-Opt. InterAb enables researchers to predict antibody specificity and antibody-antigen binding affinity. *In silico* experiments demonstrate that InterAb exhibits outstanding performance in both specificity and affinity tasks compared to current methods. InterAb-Opt is a novel antibody optimization framework guided by functional interactions, which has successfully optimized broadly neutralizing antibodies against wild-type SARS-CoV-2 and its five variants. This methodology holds considerable potential to tackle crucial challenges in immune evasion and vaccine design.

## 4 Method

### 4.1 Dataset construction

For antibody specificity prediction task, a novel dataset was constructed based on the SAbDab [43] and Observed Antibody Space (OAS) [44, 45] databases. From the SAbDab database released on January 10, 2025, 9554 antibody-antigen complexes were obtained, followed by the removal of data containing unknown amino acids or sequences longer than 1,022 residues due to limitation of computational resource. Antibody sequences with a similarity exceeding 95% were excluded using CD-Hit [46], leading to 2540 antibody-antigen binding entries. For each binding entry, two antibody sequences exhibiting a CDR-H3 sequence identity strictly below 70% with respect to the positive sample antibody were randomly sampled from the OAS database. These sequences were then paired with the corresponding antigen to construct two non-binding entries, yielding a specificity dataset (SPE7620) comprising 7,620 binding and non-binding entries in total. Among these data, two data partitioning strategies were employed: random splitting and antigen-based splitting. Under the random scheme, the dataset was randomly divided into a training set (80%), a validation set (10%), and an independent test set (10%, SPE-Test). Under the antigen-based scheme, antigens included in the training and validation sets were strictly excluded from the test set. To screen for antibodies that specifically recognize influenza A, we constructed a new dataset named FluA-Data. From previous literatures[47–56], we collected a total of 150 experimentally validated specific binding pairs between influenza A virus and antibodies. For each positive antibody sequence, we randomly selected two antibody sequences from the OAS database to serve as negative samples, resulting in a final dataset comprising 450 entries. Similarly, negative sample antibodies were required to exhibit a CDR-H3 sequence identity strictly below 70% with respect to the cognate positive sample antibody.

For antibody-antigen binding affinity prediction task, we constructed a novel dataset using SAbDab and BioMap [57] datasets. The SAbDab dataset comprises 739 entries with binding free energy (Delta G), of which 448 entries that involve antigens were selected. The BioMap dataset was derived from the 2021 Global Antibody Affinity Prediction Challenge organized by BioMap, comprising a total of 1,706 entries with Delta G annotations from 473 complexes. The data obtained from SAbDab and BioMap datasets were combined and filtered for duplicates, resulting in 1,728 entries (AFF1728). To evaluate the model’s ability to distinguish affinity differences among closely related variants, AFF1728 was randomly partitioned at the entry level, with 80% of the data used for training, 10% for validation, and the remaining 10% as a held-out test set (AFF-Test).We also constructed affinity datasets primarily focusing on SARS-CoV-2 and mutation data. By integrating our in-house experimental data with publicly available datasets[58–67], we obtained receptor binding domain (RBD) data for the wild type of SARS-CoV-2, 12 variants, and two closely related coronaviruses (Pangolin-GD and RaTG13), comprising 11,150 entries annotated with half maximal inhibitory concentration (*IC*_50_) and 1,293 entries annotated with dissociation constant (*K*_*d*_). The 12 variants mentioned above are BA.1, BA.1.1, BA.2, BA.2.12.1, BA.2.13, BA.2.75, BA.3, BA.4/5, BA.5, BQ.1.1, Beta, and Delta, respectively. For the data corresponding to each antigen, we randomly selected 80% for training, 10% for validation, and 10% as the test set. To explore the predictive capability of the model on mutations, we additionally constructed a new mutation dataset based on the SKEMPI 2.0 [36] database. A total of 7,085 entries were downloaded from the SKEMPI 2.0 database (released on January 1, 2025), from which antibody-antigen affinity data were extracted, yielding 426 mutation entries labeled as *K*_*d*_ (SKE426). For the aforementioned datasets, monomer and complex structures were predicted using ESMFold and Chai-1, respectively.

### 4.2 The architecture of the model

InterAb-Opt is an interaction-guided optimization framework (Fig. 1), in which InterAb enables precise characterization of antibody–antigen interaction. InterAb comprises three modules (Fig. 1c), namely the Inner-chain sequence module, Inner-chain structure module, and Inter-chain module. The Inner-chain sequence module utilizes pre-trained language models designed for antibodies and ESM2 to extract sequence characteristics from the sequences, subsequently integrating these into sequence embeddings. Based on the monomer structures, Inner-chain structure module extracts monomer structural features through geometric graph learning, which are then incorporated into monomer embeddings. The Inner-chain sequence and Inner-chain structure modules primarily extract features within protein chains, while the Inter-chain module is developed to extract inter-chain interaction information (complex embeddings). Utilizing the antibody-antigen complexes, AtomInter is employed to extract the interface area and further derive fine-grained features from the structure through all-atom modeling. Combining sequence embeddings, monomer embeddings, and complex embeddings, a specially designed convolutional neural network is employed to predict the binding specificity and affinity between antibodies and antigens.

### 4.3 Inner-chain sequence module

In Inner-chain sequence module (Fig. 1d), antibody representations and antigen representations are acquired using pre-trained antibody language models and ESM2 model. Given the unique characteristics of antibody data, we pre-trained specialized language models on antibody sequences, as general models are inadequate for capturing its features. RoFormer [68] was pre-trained separately on antibody heavy chain and light chain sequences.

#### 4.3.1 RoFormer

RoFormer employs an encoder-based architecture inspired by BERT, enhancing the traditional BERT framework through the incorporation of Rotary Position Embedding (RoPE). In contrast to the absolute positional encodings implemented in traditional BERT, RoPE calculates positional embeddings through rotation matrices based on relative positions, thereby enhancing the model’s ability to capture long-range dependencies within sequences. The structure of RoFormer is explained in detail as follows.

Let 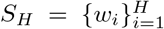 represent a sequence consisting of *H* input tokens, with *w*_*i*_ identified as the *i*th element of the sequence. The word embedding associated with *S*_*H*_ is represented as 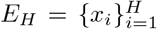, with *x*_*i*_ ∈ *R*^*d*^ signifying the d-dimensional word embedding vector for the token *w*_*i*_, excluding any positional information. The self-attention mechanism initially integrates positional information into word embeddings, converting them into representations of *q, k*, and *v*.

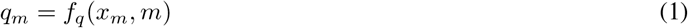

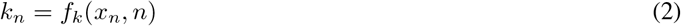

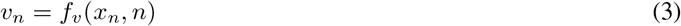

The *m*^*th*^ and *n*^*th*^ positions are integrated through *q*_*m*_, *k*_*n*_, and *v*_*n*_ with *f*_*q*_, *f*_*k*_, and *f*_*v*_. The attention weights are subsequently calculated using the *q* and *k*, while the output is determined as a weighted sum of the value representations.

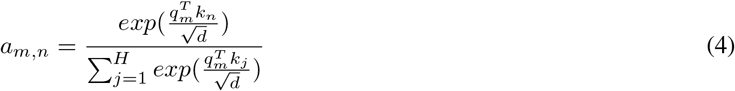

To integrate relative positional information, RoPE defines the inner product of the query *q*_*m*_ and the key *k*_*n*_ as a function *g* that accepts solely the word embeddings *x*_*m*_, *x*_*n*_ and their relative locations *m* − *n* as input.

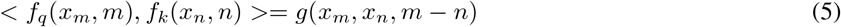

It was assumed that positional information is encoded solely in a relative format [68]. The representation of RoPE was derived as follows:

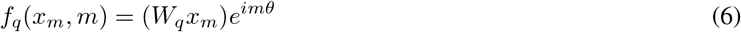

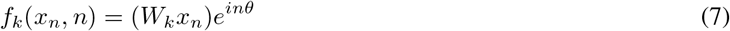

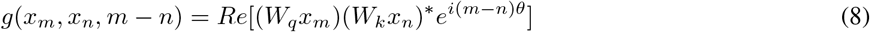

Where *Re*[·] and *θ* ∈ *R* represent the real part of a complex number and a preset non-zero constant, while (*W*_*k*_*x*_*n*_)^∗^ indicates the conjugate complex number of *W*_*k*_*x*_*n*_.

The RoPE utilizes absolute position encoding to establish relative position encoding, thus circumventing the necessity to manipulate the attention matrix. In light of the folding conformation of proteins, relative position coding may prove more effective for constructing linkage patterns among proteins. Therefore, we utilized RoFormer to pre-train antibody language models tailored for antibody data.

#### 4.3.2 Pre-trained antibody language models

Considering the significant impact of single-point mutations on antibodies, we employed a Unique Amino Acid (UAA) Tokenizer during the pre-training process. For each amino acid in antibody sequences, the UAA Tokenizer assigns a unique token along with a corresponding unique integer value. Thereafter, the integer value of each amino acid is transformed into a dense vector representation through a learnable embedding layer, enabling the model to capture positional information for each amino acid.

The Mask Amino Acid (MAA) task was utilized as a self-supervised pre-training method. Initially, a random subset of the input protein sequence is masked, followed by the prediction of these masked tokens. This approach enables pre-training models on an abundance of unlabeled data, thereby addressing the issue of limited availability of antibody data with binding specificity or affinity annotations. In practice, we randomly mask 15% of the tokens for pre-training.

During training, the intermediate size, hidden size, hidden layers, and attention heads were set to 1536, 768, 12, and 12, respectively. The AdamW optimizer was employed with a learning rate of 5 × 10^−5^. On the OAS-unpaired dataset, the heavy RoFormer model was trained on 1.3 billion antibody heavy chain sequences, while the light RoFormer model was trained on 211 million antibody light chain sequences. The final model comprises 114 million parameters. More details of the model can be seen in Supplementary Table S10.

#### 4.3.3 Pre-trained antigen ESM2 model

For antigen sequences, the pre-trained ESM2 [23] model (ESM2-150M) was utilized to extract sequence features. The ESM2 model was pre-trained using UniRef50 [69] protein sequences, based on the recognition that language models are capable of identifying evolutionary patterns within millions of sequences. Leveraging the ESM2 model’s robust expressive capabilities for general proteins, we employed it to generate sequence embeddings from antigen sequences, which were then used to predict antibody binding specificity and affinity.

### 4.4 Inner-chain structure module

The Inner-chain structure module (Fig. 1e) is employed to extract information from the structures of antibody and antigen monomers, thereby addressing the neglect of this aspect in current structure-based methods. Utilizing the monomer structures, we initially extract the structural features of the protein chains and subsequently derive monomer embeddings through geometric graph learning.

#### 4.4.1 Feature Extraction

Based on the distances between the *C*_*α*_ atoms of residues, we constructed a radius graph with a radius of 10 Å. For residue *i*, we defined vectors *u*_*i*_, *t*_*i*_, *b*_*i*_, and *s*_*i*_ as follows:

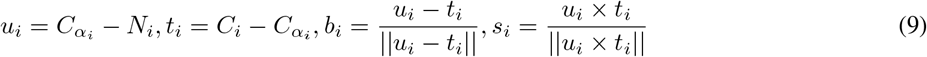

The 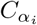, *N*_*i*_, and *C*_*i*_ are atoms of residue *i*. Based on *b*_*i*_, and *s*_*i*_, a local coordinate system *Q*_*i*_ = [*b*_*i*_, *s*_*i*_, *b*_*i*_ × *s*_*i*_] for residue *i* was established. Using this local coordinate system, we extracted node and edge features.

Node features. We constructed distance, direction, and angle features for each node. For residue *i*, we consider two atoms 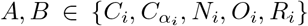, where *R*_*i*_ denotes the centroid of the sidechain atoms, and the rest are the other four atoms. The distance features are represented by a radial basis function *RBF* as *RBF* (||*A* − *B*||), *A* ≠ *B*. The direction features represent the direction of the remaining atoms relative to 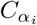 of residue *i* and are denoted as 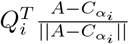. The angle features consist of the sine and cosine values of the torsion angles (*ϕ*_*i*_, *ψ*_*i*_, *ω*_*i*_). To augment the node features, ESM2 model was employed to obtain embeddings from the protein sequences, while DSSP is used to obtain structural properties.

Edge features. For atoms 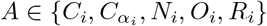 and 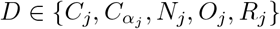 of residues *i* and *j*, we define their distance, direction, and orientation features. The distance features of residues *i* and *j* are represented as *RBF* (||*A*−*D*||), where *RBF* denotes the radial basis function. The direction features are 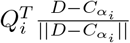, representing the direction of residue *j* relative to residue *i*. The orientation features are defined as 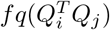, indicating the relative rotation between the local coordinate systems, where *fq* denotes a quaternion encoding function [70].

#### 4.4.2 Geometric graph learning

After obtaining the features, the geometric graph learning was used to learn monomer embeddings with several GNN layers. In order to learn structural information across multiple levels, node updates, edge updates, and global context attention are utilized during the training process.

Node updates. Define the feature vectors of node *i* and edge *j* → *i* in layer *l* as 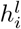 and 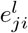, and transform them into *r*-dimensional space. The message passing operation for updating node *i* in layer *l* is described below:

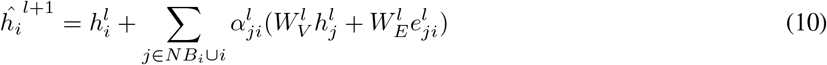

the attention weight 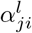 is calculated as follows:

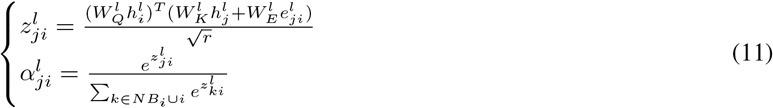

The 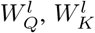, and 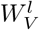 are three weight matrices used for the conversion of query, key, and value representations. The edge vectors were utilized to enhance the key and value representations through weight matrices 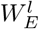, while *NB*_*i*_ denotes the neighbors of node *i*.

Edge update. Edge features are updated based on neighboring nodes to improve the representation of interactions between residues.

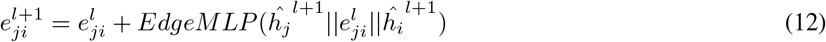

Where || indicates the concatenation operation and *EdgeMLP* represents the MLP operation conducted on edges.

Global context attention. While applying global self-attention across the entire protein can capture the global features, its substantial computational cost restricts its practical utility. To address this issue, we compute a global context vector before carrying out gate attention operations, with reference to PiFold [27].

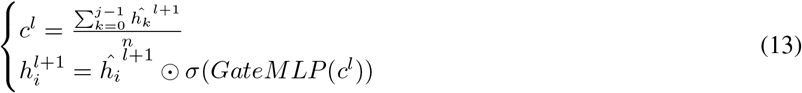

Here, *j* symbolizes the number of residues in a protein, *σ* denotes the sigmoid function, is the element-wise product operation, and *GateMLP* represents the MLP designed for gated attention.

Through geometric graph learning, we obtained monomer embeddings from the monomer structures of antibodies and antigens. These embeddings capture critical information within the structures and provide insights into the structural details prior to the binding of antibodies and antigens.

### 4.5 Inter-chain module (AtomInter)

We propose an all-atom modeling module, AtomInter, to extract complex embeddings from antibody-antigen complexes (Fig. 1f). To capture essential information regarding the antibody-antigen interactions and reduce memory consumption, we delimited the antibody-antigen interface area based on *C*_*α*_ distance metrics (10 Å). Utilizing the interface area, we extracted atomic structural features from the atomic sequences along with their corresponding coordinates and positional information.

The atomic sequence is encoded through one-hot encoding based on atomic type, resulting in a 37-dimensional atomic sequence feature vector. The three-dimensional coordinates of each substructure were normalized to address inconsistent structural coordinates, thereby enhancing the atomic sequence features. Finally, by integrating the position encodings for the atomic sequences and the corresponding amino acid positions, we obtained a total of 42-dimensional atomic structural features *Fa*.

For each atom, the updated atomic features *Fa*^*l*+1^ in layer *l* + 1 is obtained from the features *Fa*^*l*^ in layer *l* as follows:

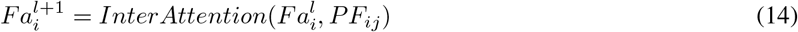

The *PF*_*ij*_ represents pairwise atomic structure features. It is calculated by first computing the atom-level pairwise distances. Then, these distances are integrated with the atoms’ corresponding reference space through the use of atomic coordinates. Finally, it serves as a bias in the window attention mechanism, as referenced in AlphaFold 3.

The *InterAttention* is defined as follows. Initially, the input projections are performed on *Fa*_*i*_. For each attention head *h*_*a*_ (ranging from 1 to *N*_head_), we obtain the query vector 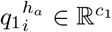, key vector 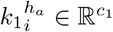, and value vector 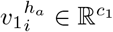. Subsequently, the attention computation is executed. The attention score 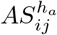 for each attention head *h*_*a*_ is computed as follows:

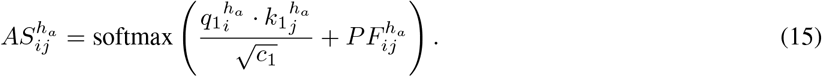

This involves taking the dot product of the transpose of 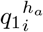 and 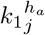, scaling it by 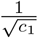, adding the pair bias 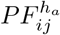, and applying the softmax function for normalization. After normalization by softmax, the sum of all 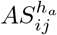 equals 1, which facilitates subsequent calculations. Then, the node feature *Fa*_*i*_ is updated. For each attention head *h*_*a*_, the weighted sum of 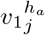 by attention scores is calculated. The updated feature is given by

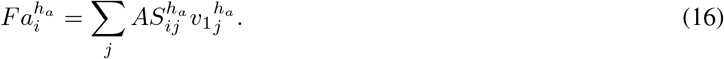

The results from different attention heads *h*_*a*_ are concatenated to obtain the updated *Fa*_*i*_, enabling *Fa*_*i*_ to learn the feature information of other atoms. Finally, the *InterAttention* method returns the updated set of *Fa*_*i*_ (i.e., the previously defined 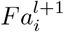), which effectively incorporates pair bias information during the update of atomic features. This enhances the learning of interactions between atoms and provides richer feature representations.

Additionally, to avoid the initial features from being overlooked, we employ residual connections to enhance atomic attention.

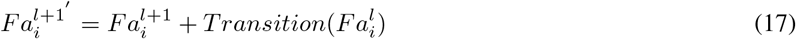

Where *Transition* refers to a SwiGLU [71] transition block with adaptive LayerNorm. Then, a stacked CNN model is utilized to extract complex embeddings *C*_*emb*_ from 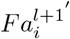.

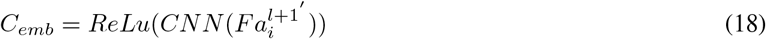

### 4.6 InterAb-Opt

InterAb-Opt was proposed to optimize broadly neutralizing antibodies based on the functional insights provided by InterAb. Employing the antibody sequence and antibody-antigen structure, we first identified the critical sites of the antibody (Fig. 4a). Subsequently, our previously developed method, IgGM [30], was utilized to generate candidate antibodies, which were then screened using InterAb to identify those demonstrating broad-spectrum activity.

Identifying critical sites in antibodies is a major challenge in antibody optimization, and we have determined the critical sites in phases. Initially, we selected residues on the antibody within an 8Å distance from the antigen as candidate sites (25 residues for R1-32). Subsequently, *in silico* alanine scanning via InterAb was exploited to determine the critical sites (13 residues for R1-32) from the candidate sites. The *in silico* alanine scanning process (Fig. 4b) involved mutating each candidate residue to alanine, followed by predicting the binding affinity of the mutated antibody using InterAb. The residues were then ranked based on the degree of affinity reduction, and the top 50% were designated as critical sites. Particularly, if the original residue is alanine, we consider the site as a critical site. Fig. 4c illustrates the 13 critical sites selected for R1-32.

Upon identifying the critical sites, we utilized IgGM to generate 1,000 antibodies for each antigen. Using these generated antibodies, a candidate antibody library was constructed, and InterAb was utilized to assess the binding affinities between the candidate antibodies and all antigens. The average binding affinity of the antibodies against these antigens was calculated, and antibodies with high affinity were selected for experimental validation through biolayer interferometry (BLI, details shown in Section 4.12).

### 4.7 Transforming *K*_*d*_ and *IC*_50_ values into log-space

The dissociation constant (*K*_*d*_) is used to quantify binding affinity in the SARS-CoV-2 and SKE426 datasets. However, due to the considerable variation in the original *K*_*d*_ values, we transform the *K*_*d*_ values into log space to achieve a more uniform distribution of the data.

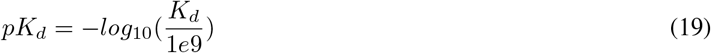

The *pK*_*d*_ represents the transformed value, which reduces the data range compared to the original *K*_*d*_ values.

Similarly, we transformed the *IC*_50_ values into the log space as *log*_10_(*IC*_50_):

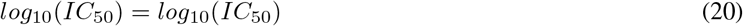

### 4.8 Predicting monomer structures by ESMFold

ESMFold [23] was used to predict monomer structures. Given the premise that language models can effectively capture evolutionary patterns from extensive sequences, a large language model (ESMFold) comprising up to 15 billion parameters is proposed. By replacing multiple-sequence alignments (MSAs) with a language model, ESMFold achieves a speed nearly 60 times faster than AlphaFold2, enabling the first evolutionary-scale structural characterization of a metagenomic resource. The monomer structures of antibodies and antigens can be rapidly predicted using ESMFold.

### 4.9 Predicting complex structures by Chai-1

Chai-1 was used to predict antibody-antigen complex structures. Chai-1 is a multi-modal foundation model designed for predicting molecular structures, demonstrating superior performance across various tasks such as protein multimer prediction and protein-ligand structure prediction. Experimental constraints, such as those derived from epitope mapping, can be leveraged to enhance the accuracy of predictions. Furthermore, it can deliver rapid and accurate predictions without MSAs in single sequence mode, making it suitable for high-throughput applications. For each antibody-antigen pair, we generated five complex structures using Chai-1 and selected the structure with the highest ipTM score as the final predicted structure.

### 4.10 Expression and purification of optimized antibodies

The heavy chain (IgH) and light chain (IgL) genes of optimized antibodies were amplified and cloned into the expression vector pCMV3. Expi293F cells (Thermo Fisher Scientific, A14527) were cultured in Expi293F culture medium (Thermo Fisher Scientific, A1435101) at 37 °C with 8% CO2 for protein expression. When the density of Expi293F cells reached 2.5 × 10^6^ cells/mL, the paired IgH and IgL plasmids were transiently co-transfected into the Expi293F cells at a ratio of 1:1 using the transfection reagent polyethylenimine (PEI). Five days post-transfection at 33 °C, supernatants were harvested by centrifugation and incubated with Protein A Resin (Genscript, China) at room temperature for 2 h to enable antibody binding. After the resin was washed with PBS (Gibco, pH 7.4), the antibodies bound to the resin were eluted using 0.1 M citric acid (pH 3.0), and neutralized immediately with an equal volume of 1 M Tris-HCl (pH 8.0), to maintain the pH within the neutral range. Subsequently, fractions containing antibodies were concentrated and buffer-exchanged into PBS using a 50 kDa MWCO Amicon Ultra filtration (Merck Millipore) at 4 °C. Purified antibodies were aliquoted, flash-frozen, and stored at −80 °C until needed.

### 4.11 Expression and purification of SARS-CoV-2 RBD

The coding sequence of SARS-CoV-2 RBD (residues 332–527), fused with an N-terminal *µ*-phosphatase signal peptide and a C-terminal hexahistidine tag, was cloned into the expression vector pcDNA3.1. Similar to antibody production, the RBD plasmid was transiently transfected into Expi293F cells using the transfection reagent polyethylenimine (PEI) when the cell density reached 2.5 × 10^6^ cells/mL. Five days post-transfection at 33 °C, the culture supernatant was collected by centrifugation and supplemented with 25 mM phosphate, pH 8.0, 300 mM NaCl, and 5 mM imidazole, and recirculated onto a HiTrap TALON crude column (Cytiva). Subsequently, the column was washed with buffer A (25 mM phosphate, pH 8.0, 300 mM NaCl, 5 mM imidazole), and the RBD protein was eluted with a 100 mL linear gradient to 100% buffer B (25 mM phosphate, pH 8.0, 300 mM NaCl, 500 mM imidazole). Fractions containing the target protein were pooled, concentrated with a 10 kDa MWCO Amicon Ultra filtration (Merck Millipore). Eventually, the RBD protein was further purified and buffer-exchanged into PBS using a Superdex 200 increase 10/300 GL column (Cytiva). All SARS-CoV-2 RBD variants were expressed and purified following the aforementioned protocol. Purified RBDs were aliquoted, flash frozen, and stored at −80 °C until needed.

### 4.12 Biolayer interferometry (BLI) binding assay

Binding kinetics and affinities of optimized antibodies against the RBDs of wild-type SARS-CoV-2 and the variants were assessed by biolayer interferometry (BLI) on an Octet-RED96 (ForteBio). All experimental procedures were performed at 25 °C and all proteins were diluted to PBST buffer (PBS with 1 mg/mL BSA and 0.02% v/v Tween-20). Initially, antibodies (11 µg/mL) were immobilized onto Protein A biosensors. After a 60 s baseline step in PBST buffer, biosensors loaded with antibodies were exposed (300 s) to various RBDs (200 nM) to measure association. Subsequently, the biosensors were dipped into PBST buffer (600 s) to measure dissociation of RBDs from the biosensor surface. A blank reference was set for each reaction. Data was reference-subtracted and analyzed using the Octet Analysis Studio software v.12.2, fitting the data to 1:1 or 2:1 binding model to determine kinetic parameters. Results were plotted using GraphPad Prism 8.0.

### 4.13 Neutralization assay

The neutralizing activity of antibodies targeting wild-type SARS-CoV-2 and several variants (including Lambda, BQ.1.1, and EG.5.1) was assessed using a pseudovirus-based neutralization assay according to previously described methods [72]. Antibody samples were initially diluted to 10 *µ*g/mL and subjected to eight gradient dilutions (3× serial dilutions). The pseudovirus was adjusted to a concentration of 1.3 × 10^4^ TCID_50_/mL. Following a 1-hour incubation of the antibody-virus mixture at 37^°^C, HEK293T-ACE2 cells were introduced. After 24 hours of culture, a luciferase detection reagent was added, and luminescence signals were measured using a multimode microplate reader (PE EnSight). Data analysis and *IC*_50_ calculations were performed with GraphPad Prism 8.0.

### 4.14 Implementation and evaluation

The models were trained on antibody-antigen binding specificity and affinity datasets, and their performance was evaluated on multiple test sets. In the training phase, the binary cross-entropy loss was utilized for the specificity task, while mean squared error loss was applied to the affinity task. The Adam optimizer with a learning rate of 1 × 10^−4^ was employed to optimize the model, with a maximum of 100 epochs set for the training process. Early stopping with a patience of 5 was utilized, while a dropout rate of 0.2 was applied to prevent overfitting. For specificity task, the trained model was evaluated on the independent test set SPE-Test. For affinity task, the model was assessed on the independent test set AFF-Test, as well as the SARS-CoV-2 test set and the mutation test set. To comprehensively evaluate the model performance, we employed a range of metrics, including area under the receiver operating characteristic curve (AUC), area under the precision-recall curve (AUPR), recall, precision, F1-score (F1), Matthews correlation coefficient (MCC), Accuracy (ACC), Pearson’s correlation, Spearman’s correlation, Root mean square error (RMSE), and Mean absolute error (MAE). The methodologies for calculating these metrics are detailed in the Supplementary information (Evaluation metrics).

## 5 Data Availability

The dataset concerning antibody specificity, SPE7620, is available on GitHub (https://github.com/YidongSong/InterAb/tree/main/data/SPE7620.csv). The dataset AFF1728, which includes information on antibody-antigen affinity compiled from the SAbDab database and the BioMap dataset, can be found at (https://github.com/YidongSong/InterAb/tree/main/data/AFF1728.csv). The mutation dataset compiled from the SKEMPI 2.0 database is available at (https://github.com/YidongSong/InterAb/tree/main/data/SKE426.csv).

## 6 Code Availability

The source code of InterAb is available at https://github.com/YidongSong/InterAb. The pre-trained antibody models, ESM2 model, and the trained InterAb model can be downloaded from Zenodo (https://zenodo.org/records/21785933).

## 7 Acknowledgements

This work was supported by the National Natural Science Foundation of China (12426303 to W.Z.)

